# Multi-ContrastiveVAE disentangles perturbation effects in single cell images from optical pooled screens

**DOI:** 10.1101/2023.11.28.569094

**Authors:** Zitong Jerry Wang, Romain Lopez, Jan-Christian Hütter, Takamasa Kudo, Heming Yao, Philipp Hanslovsky, Burkhard Höckendorf, Rahul Moran, David Richmond, Aviv Regev

**Author notes:** Work performed during an internship at Genentech, Inc (Zitong Jerry Wang).

## Abstract

Optical pooled screens (OPS) enable comprehensive and cost-effective interrogation of gene function by measuring microscopy images of millions of cells across thousands of perturbations. However, the analysis of OPS data still mainly relies on hand-crafted features, even though these are difficult to deploy across complex data sets. This is because most unsupervised feature extraction methods based on neural networks (such as auto-encoders) have difficulty isolating the effect of perturbations from the natural variations across cells and experimental batches. Here, we propose a contrastive analysis framework that can more effectively disentangle the phenotypes caused by perturbation from natural cell-cell heterogeneity present in an unperturbed cell population. We demonstrate this approach by analyzing a large data set of over 30 million cells imaged across more than 5, 000 genetic perturbations, showing that our method significantly outperforms traditional approaches in generating biologically-informative embeddings and mitigating technical artifacts. Furthermore, the interpretable part of our model distinguishes perturbations that generate novel phenotypes from the ones that only shift the distribution of existing phenotypes. Our approach can be readily applied to other small-molecule and genetic perturbation data sets with highly multiplexed images, enhancing the efficiency and precision in identifying and interpreting perturbation-specific phenotypic patterns, paving the way for deeper insights and discoveries in OPS analysis.

## 1 Introduction

Large-scale, pooled genetic perturbation cell screens with high content readout enable systematic interrogation of gene function [1]. Among these, optical pooled screens (OPS) that employ cell imaging as phenotypic readout for characterizing perturbation effects are particularly compelling, because they offer high-throughput, complex readout at low-cost [2].

Traditional phenotype analysis in OPS and other image-based screens requires extracting from each image a set of hand-crafted morphological features [3, 4]. Based on these, established tools like CellProfiler (CP) usually apply a set of pre-defined filters to images of individual cells in order to summarize the data from each single cell into a data point using thousands of features. There are several limitations to these approaches. First, because the filters have been engineered on previous data sets, with different cell types and experimental conditions, hand-crafted features may be inflexible to capture novel morphological phenotypes. Indeed, because cellular morphology drastically changes across contexts and biological systems, the set of optimal features may be different for each experiment. Second, those extracted feature are limited in their ability to capture interactions between channels. For example, CP accomplishes this by applying filters to pairs of channels (usually for images with two to five channels). However, this will be difficult to scale in order to capture more complex patterns in images with tens to hundreds of channels which are becoming increasingly common [4].

Advances in deep learning could overcome the limitations of hand-crafted features by learning representations directly from data [3, 5]. In particular, the Variational Auto-Encoder (VAE) is a powerful deep generative framework to capture latent structure in complex data distributions in an unsupervised manner [6, 7]. However, in the context of learning perturbation effects from images of cells, standard VAE implementations suffer from the difficulty of isolating perturbation effects from natural cell-cell heterogeneity (e.g., due to stages of the cell cycle), which can exhibit much greater variation compared to the phenotypic effect of perturbations.

Contrastive analysis (CA) offers a potential solution to identify and isolate patterns induced by perturbations using a background data set to remove those natural variations [8]. In our setting, the background data set is composed of images of control cells that have not been perturbed. For example, contrastive principal components analysis (cPCA) seeks to identify salient principal components in a target data set by identifying linear combinations of features that are enriched in that data set relative to a background data set, rather than those that simply have the most variance [9]. Recently, CA methods based on neural networks have proven effective in discovering nonlinear latent features that are enriched in one dataset compared to another [10–14]. These approaches often assume that only two datasets are being processed, both stemming from identically and independently distributed data distributions: the background distribution and the target distribution. This approach is limited in that it does not explicitly model each perturbation as a unique distribution for comparison with the background distribution.

Here, we propose Multi-ContrastiveVAE, a CA framework for the setting of comparing multiple data sets to a reference data set, with a specific architecture tailored for cell imaging data sets from OPS. We applied Multi-ContrastiveVAE to a large-scale imaging dataset with over 30 million cell images across more than 5, 000 genetic perturbations, as detailed in [15]. Our approach more accurately identifies perturbation-specific phenotypes compared to non-contrastive methods, effectively distinguishing them from cell-to-cell variations that persist across perturbations. Multi-ContrastiveVAE effectively separates multiple sources of technical artifacts from single-cell images, including non-biological variations due to batch effects, uneven plating, and uneven field of view (FOV) illumination. Furthermore, Multi-ContrastiveVAE disentangles perturbation effects into separate latent spaces depending on whether the perturbation induces novel phenotypes unseen in the control cell population. Multi-ContrastiveVAE is readily applicable to other perturbation data sets with highly multiplexed images, including both drug and genetic perturbation.

## 2 Methods

We developed the Multi-Contrastive Variational Autoencoder (mcVAE) to disentangle perturbation effects in large-scale perturbation data sets from natural and technical cell-to-cell variations. This model extends the CA framework, by allowing for the comparison of multiple groups against a single reference group (the background data set composed of control, unperturbed cells). Our framework is based on the generative model illustrated in Figure 1A, and builds on recent work on CA [10, 14].

**Figure 1:**
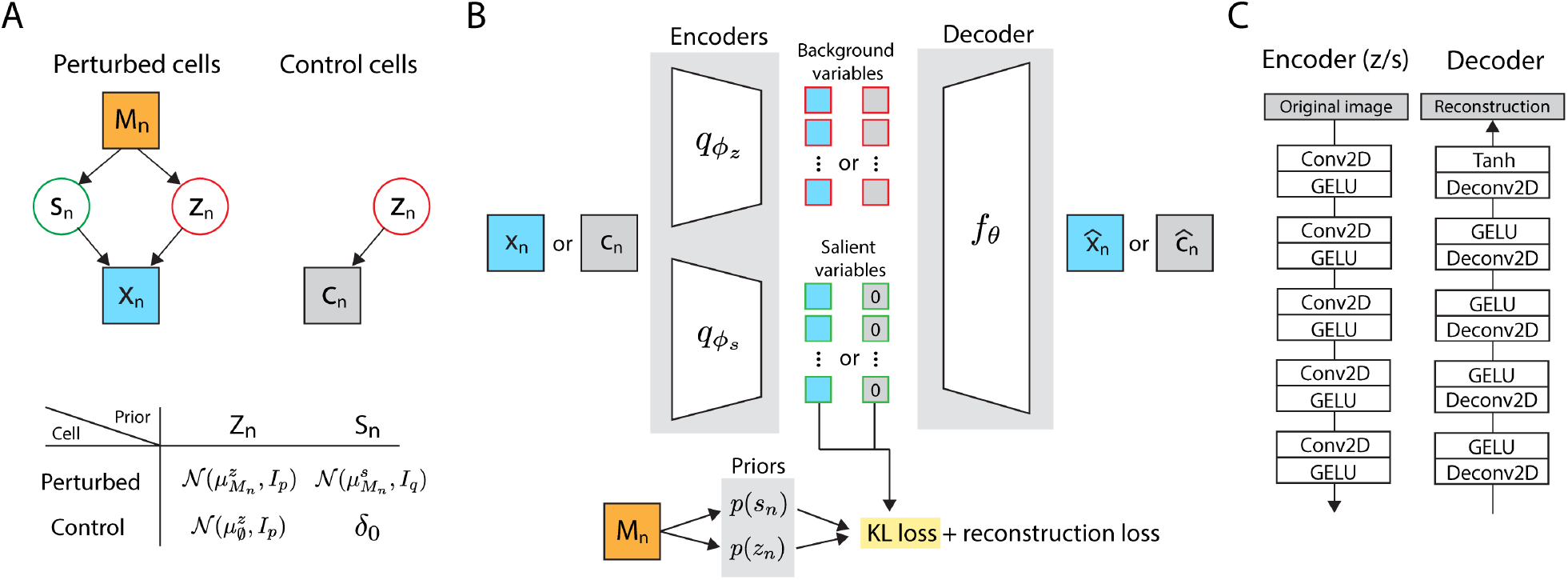
A contrastive analysis framework for analyzing single cell images from optical pooled screens. (A) Generative model for perturbed and control cells. The table shows the prior distributions used for the background and salient variables of the perturbed and control populations. (B) Schematic of the Multi-Contrastive variational autoencoder (mcVAE) framework. (C) Neural network architectures of the encoder and the decoder.

### Generative Model

For each cell *n*, we observe the perturbation label *M*_*n*_ ∈ {∅, 1, …, *K*}, where ∅ denotes a non-targeting control (NTC) perturbation (i.e., no effect) and *K* denotes the number of distinct perturbations. Let *n* be for now a perturbed cell, that is *M*_*n*_ ≠ ∅. Let latent variable

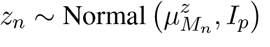

be a low-dimensional random vector encoding cellular variations that naturally exist in the control cells (the background data set). The mean of the prior *p*(*z*_*n*_ | *M*_*n*_) varies with the perturbation label *M*_*n*_ to account for the fact that perturbations may shift the density of cells towards certain preexisting cell states from the control population. We refer to *z* as the *background* latent space, or embedding. Then, let latent variable

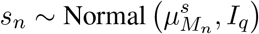

be a low-dimensional vector encoding variations due to perturbations. The mean of the prior *p*(*s*_*n*_ | *M*_*n*_) is shifted by 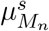 to account for the fact that different perturbations may incur different changes in the data distribution. We refer to *s* as the *salient* latent space, or embedding. All the images *x*_*n*_ ∈ *ℝ*^*d*^ have the same number of pixels *d*. We assume that each pixel *j* in each image *x*_*nj*_ is generated as:

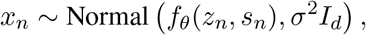

where *f*_*θ*_ is a neural network taking value in the hypercube [−1, 1]^*d*^.

In order to break symmetry between latent variables *s* and *z*, we exploit the control cells. The data distribution for the images of the control cells (i.e., *M*_*n*_ = ∅) is the result of an intervention

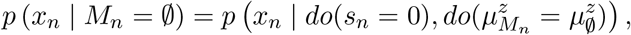

where 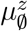 denotes the mean of the prior embedding for the embedding of the control cells (it could be set to zero without loss of generality). This assumption is classical in CA, and helps enforce the semantic that only *z* (the remaining latent variable) may be used to describe data from the control cells.

### Interpretation and Significance

Departing from established CA models, mcVAE includes additional parameters *µ*^*s*^ and *µ*^*z*^ that capture the heterogeneity of perturbations. The former captures the fact that perturbations could induce novel phenotypes (to be captured by the salient variables). The latter biases cell states after perturbation towards phenotypes that already existed in the natural population. This represents a significant conceptual departure from the original framework where the background space was interpreted as containing only uninteresting variation, while the perturbation effect resided solely within the salient space.

### Variational Inference

The marginal probability of the data *p*(*x* | *M*) is intractable. We therefore proceed to posterior approximation with variational inference to learn the model’s parameters. In particular, we use a mean-field variational distr tion:

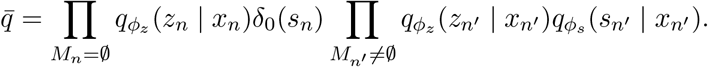

As in the VAE framework [6], each *q*_*ϕz*_ (*z* | *x*) and *q*_*ϕs*_ (*s* | *x*) follows a Gaussian distribution with a diagonal covariance matrix. We optimize a composite objective function, corresponding to the sum of the evidence lower bound (ELBO) for perturbed cells, and for control cells.

Adapting the work of [10], the ELBO for a perturbed cell *n* is derived as:

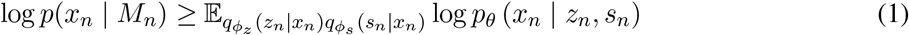

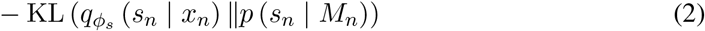

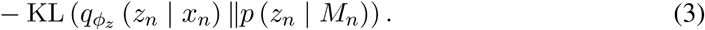

Similarly, the ELBO for a unperturbed cell *n* is derived as:

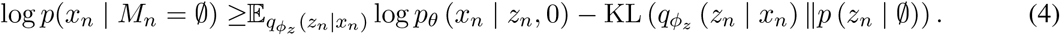

We summarized those computations in the schematic from Figure 1B.

### Architecture and Training Details

For the neural networks used in the generative model, and the amortization of the approximate posterior, we use a simple convolutional architecture with five convolutional layers for the approximate posterior, and five transpose-convolutional layers for the generative model (Figure 1C). We used the Adam optimizer [16] with a learning rate of 10^−4^. Each mini-batch was constructed to contain an equal number of perturbed and control cells. We applied the inverse hyperbolic sine transformation as well as normalization to the training images during pre-processing. All models presented in this paper were trained for 2 epochs with early stopping based on the validation loss, and the dimensions of *z* and *s* were set to 32. We regularized the model using the Wasserstein regularization [14] to encourage independence between salient and background latent features.

## 3 Results

We apply mcVAE to a public imaging data set from a large-scale CRISPR-knockout OPS that profiled 31 million HeLa cells affected by 5, 000 distinct genetic perturbations [15]. Four single guide RNA (sgRNAs) were used for each gene target, together with 250 non-targeting control (NTC) sgRNAs that do not target any gene. For 31 million cells, there is a median of around 6, 000 cells imaged per gene target across each set of four guides. To obtain single-cell images, we extracted 100 *×* 100 pixel crops without segmenting cells and perturbation label assignments for each cell (Appendix A.1). We then assessed how well the mcVAE embedding reflected known biology, how well it isolated technical artifacts from biological variation, and how the salient and background space differ in terms of the perturbation effects they capture.

### 3.1 Assessment of Embeddings Quality based on Protein Complexes

We evaluated the effectiveness of our learned embeddings in capturing known biological functions by employing them to predict established protein complexes [17]. It is well established that genetic perturbations of genes encoding different subunits of the same protein complex are more likely to yield similar cellular phenotypes. Thus, we expect the images of cells perturbed for such genes to be closer in embedding space compared with those from cells perturbed for genes genes that do not belong to the same complex. To this end, we used as ground truth the CORUM database [18], the most extensive publicly available collection of manually curated mammalian protein complexes. We set all aggregated gene embeddings up to a given cut-off as true relationships, used those to predict which gene pairs co-occur in a CORUM complex, and then evaluated the classification performance by comparing the precision-recall curve of various methods, generated by setting different distance thresholds for determining whether two genes are part of the same complex.

We compare the performance of mcVAE to three baseline models: standard VAE, contrastive VAE (cVAE), and CellProfiler (CP). Both the standard VAE and cVAE consist of the same encoder and decoder architecture (Figure 1C). The standard VAE has a single encoder and decoder, and the cVAE consists of two encoder and a shared decoder but does not use the perturbation label to adjust the priors of its latent variables. To keep the dimension of the latent space consistent across all models, instead of using all CP features, and instead we first perform PCA at the cell-level and then take the first 64 PCs (77% variance explained).

The standard VAE trailed in performance, being markedly surpassed by the cVAE (Figure 2A). Importantly, mcVAE not only exceeded the performance of the cVAE but did so to a degree comparable to the improvement seen previously going from the standard VAE to the cVAE, thereby matching the performance of CP.

**Figure 2:**
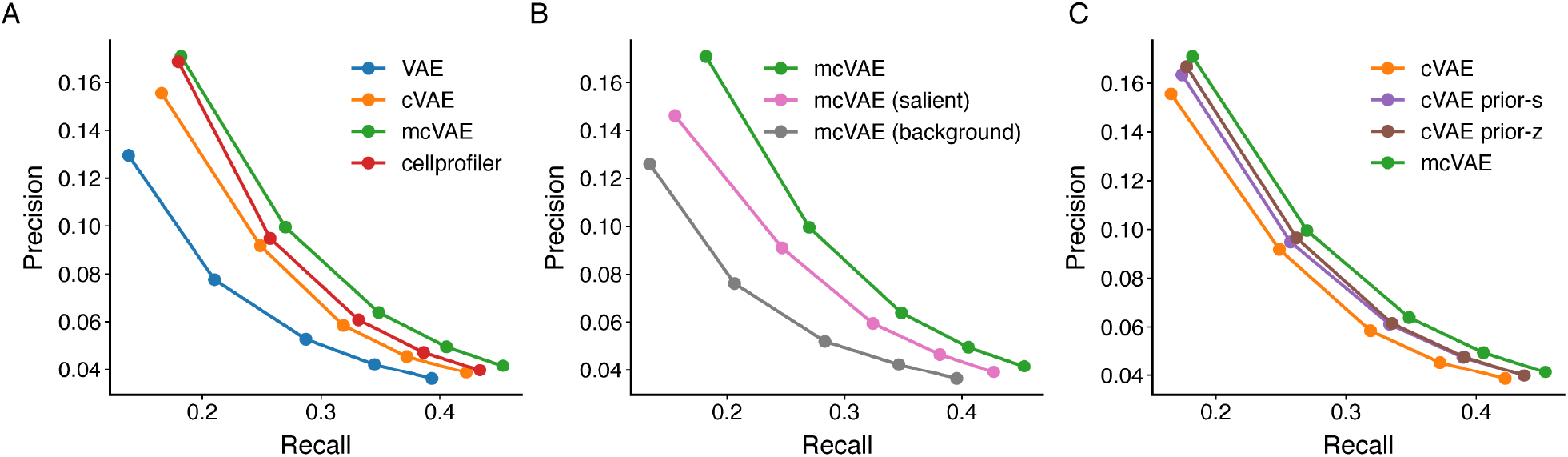
Multi-Contrastive VAE outperforms contrastiveVAE in protein complex identification. (A) Precision-recall curve of CORUM identification task for vanilla VAE, contrastiveVAE (cVAE), multi-contrastiveVAE (mcVAE) and using CP features; all four embedding spaces have 64 dimensions. (B) Precision-recall curve when using background alone, salient alone, or a concatenation of both sets of latent variable. (C) Precision-recall curve for cVAE, cVAE with perturbation label fed into just the salient prior, cVAE with perturbation label fed into just the background prior, and our mcVAE where perturbation labels are fed into both the salient and background prior

We further evaluated how the background and salient embedding of mcVAE performs separately in identifying members of the same protein complexes (Figure 2B). Interestingly, the best performance for CORUM identifiability was not achieved by discarding the background information; instead, it was attained by concatenating the salient features with the background. Thus it is not surprising that we obtain sub-optimal performance when we remove either the label information from the salient prior or the background prior (Figure 2C). These findings emphasize the critical role that both the salient and background latent variables play in identifying gene functions (defined here by protein complex membership), and highlights the strength of mcVAE in elucidating complex biological relationships.

### 3.2 Performance at Disentangling Multiple Sources of Technical Artifacts

Multi-Contrastive VAE automatically isolates multiple, intricate technical artifacts found in cell images without any prior information. First, well-to-well batch variations can emerge from multiple factors, such as subtle differences in culture (e.g. cell density) or staining conditions (Figure 3A, top). As a result, the background embeddings of cells show grouping by well in the UMAP projection, but cells from different wells are well-mixed in the salient space, indicating the salient space is nearly free of batch effect. Next, uneven illumination of a field of view can cause cells near the center to appear brighter than those near the edge. Background embedding of cells capture this variation, while the salient space appears well-mixed, indicating removal of this technical variation (Figure 3A, middle). Lastly, background cell embeddings are separated based on their position in a well, influenced by uneven cell density affecting cell shape and size, while this source of variation is again not discernible in the salient space (Figure 3A, bottom). Note that for both the position in FOV and well, the corresponding UMAPs are only showing cells from a single well to better illustrate these additional variations beyond batch effect.

**Figure 3:**
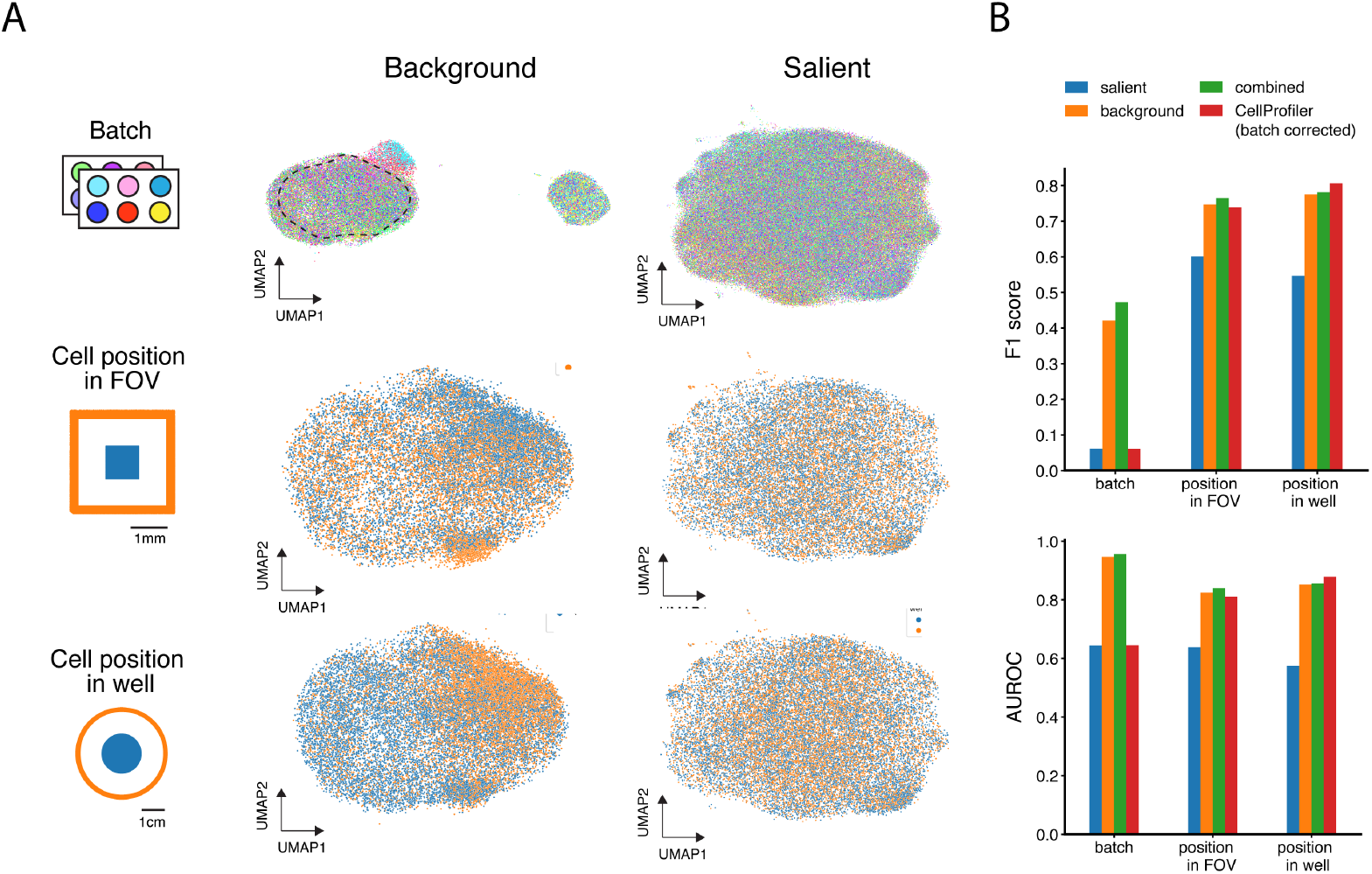
Multiple experimental technical artifacts are found in both the background space, and CellProfiler features, while the salient space is nearly free of these artifact patterns. (A) UMAP projections of background and salient embedding for individual cells colored by their batch (well), position in an image’s FOV (edge/center), and position in a well (edge/cell). The embeddings showing FOV and well position are for cells from one particular batch, which is also demarcated by dashed black line in the top left UMAP. (B) Performance metrics for a logistic regression model trained to predict the technical covariates from panel (A) using the cell embeddings from salient space, background space, salient-background concatenated, and batch-corrected CP features.

To quantify the presence of technical variation in different embedding spaces, we trained logistic regression models to predict various technical covariates from the cell embeddings (Figure 3B). The salient embeddings are mostly free of technical artifacts, as evident from having the poorest prediction performance, measured by the F1 score and area under receiver operating characteristic curve (AUROC). In contrast, both the background and CP embeddings contain significant technical variations in terms of FOV position and well position. The CP embedding we used was batch-corrected by standardizing against the NTC in the corresponding batch. While effective at removing batch effects, such correction was unable to address the other more intricate sources of technical artifacts.

### 3.3 Background and Salient Embeddings Excel at Predicting Distinct Gene Functions

Though both salient and background embeddings can accurately classify gene functions, their performance varies significantly based on the functional group. A UMAP projection of guide embeddings (cells aggregated to the level of CRISPR guides) (Figure 4A), shows that genes of the same functional groups (as assigned in [15]) tend to cluster together, suggesting that the embedding space is rich in biological information.

**Figure 4:**
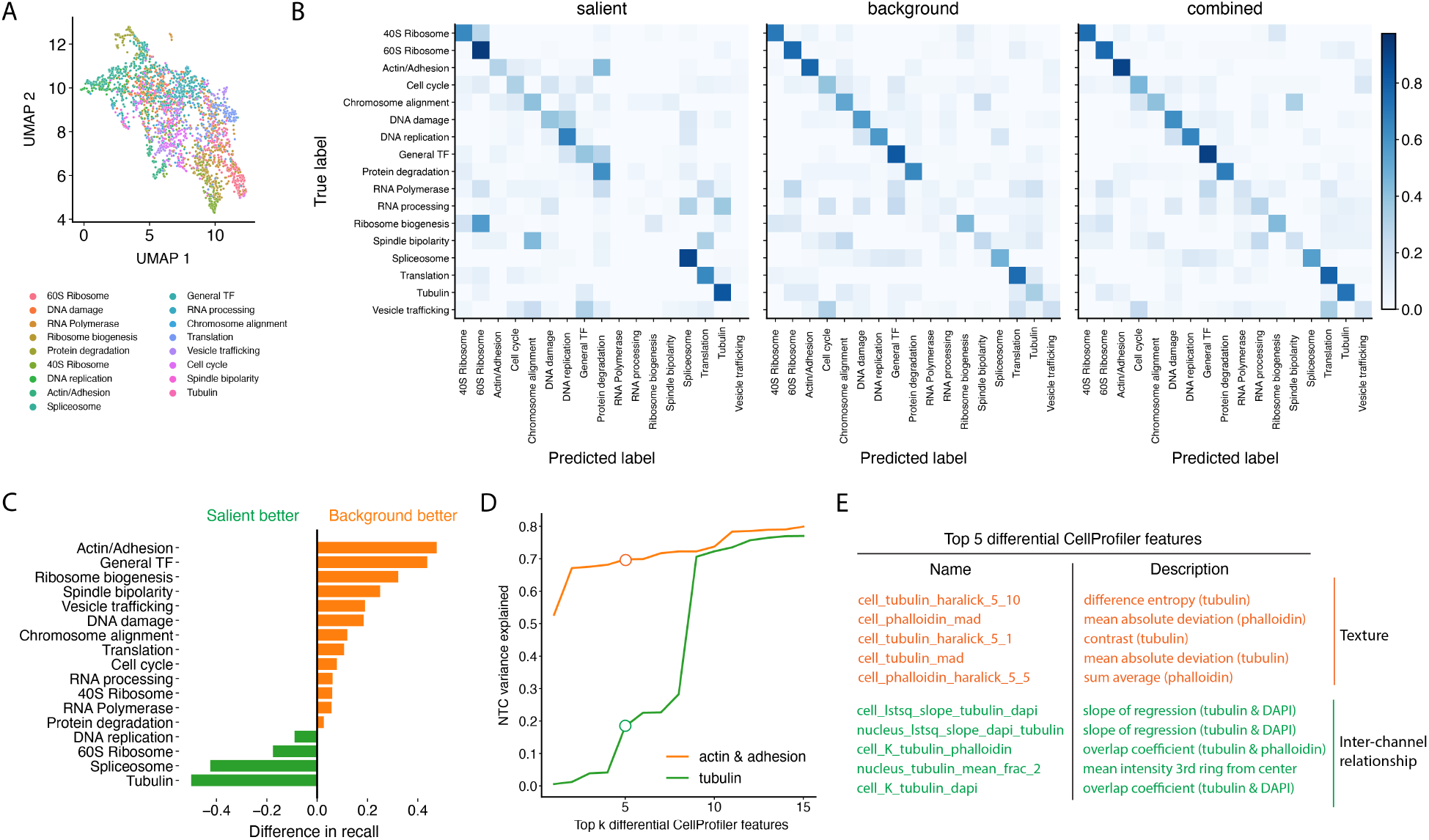
The salient space better delineates perturbations that induce novel phenotypes, while the background space highlights perturbations that shift the distribution of cells within existing phenotypes. (A) UMAP projection of salient and background cell embeddings aggregated to the perturbation level, colored by the functional group of the perturbed gene. (B) Confusion matrices for separate logistic regression models trained to classify gene function from guide embeddings in different spaces, normalized row-wise. (C) Difference in recall between model trained on salient vs background embedding. (D) Percent of variations in the first PC of NTCs explained by different number of top differentially altered features, identified by multiple hypothesis testing with Bonferroni adjustment and ranked by robust z-score against NTCs. (E) Name and description of the top five differentially altered CP features.

Salient and background spaces each excel at predicting different gene functions. The confusion matrices (Figure 4B) obtained by training logistic regression models to classify the perturbed gene into one of the functional groups from different guide-level representations, show that using the combined embedding performs best compared to using only the salient or background embedding alone.Furthermore, the difference in performance between the salient and background space vary significantly between functional groups (Figure 4B, C). The background embedding excels for predicting functional groups such as actin cytoskeleton/adhesion, spindle bipolarity, and chromosome alignment, while the salient embedding is better at predicting genes with tubulin and spliceosome annotations. The misclassifications as shown on the offdiagonal of the confusion matrices seem to associate closely related functional groups. In the salient space, for example, genes associated with ribosome biogenesis are often misclassified as part of the 60S ribosome, and similarly between genes for chromosome alignment and spindle bipolarity.

We posit that certain gene functions, such as adhesion and mitosis, were predicted better by the background embedding because perturbing these genes likely affects phenotypes, like cell size and cell cycle stage, which naturally vary within unperturbed cells (with NTC guides). As a result, the altered features are predominantly captured by the background space. Conversely, perturbing genes in the tubulin group, which is only well-predicted by the salient space, produces novel features do not appear in unperturbed cells. Indeed, using the top five altered CP features from the actin & adhesion perturbations, we can explain 70% of the variation in the first principle component (PC) of cells with NTC guides, whereas only 20% can be explained by the top five altered CP features from the tubulin perturbations (Figure 4D). Comparing the actual CP features significantly altered by perturbations to either groups, we found that perturbing the actin & adhesion group primarily affects spatial heterogeneity of a molecule, while perturbing the tubulin group mainly affects cross-correlation between different molecular species (Figure 4E). This result suggests that spatial cross-correlation between molecular structures are fairly preserved, and perturbing only tubulin disrupts these cross-correlations to generate a novel phenotype rarely seen in natural cell populations.

In summary, the differences in classification performance between the salient and background embeddings stem from the nature of the gene perturbations, specifically whether they generate new phenotypes or merely shift the distribution of existing phenotypes, shedding light on the complex interaction between gene functions and cell biological phenotypes.

## 4 Discussion

In this work, we present Multi-Contrastive VAE (mcVAE), a method for disentangling perturbation effects by comparing multiple treatment groups to a single reference/control group. We applied mcVAE to a recent largescale optical pooled screen dataset [15] consisting of over 30 millions cells spanning more than 5, 000 genetic perturbations to show that it can effectively remove technical imaging artifacts to identify perturbations that generate novel phenotypes.

Although mcVAE effectively isolated novel phenotypes in the salient space, additional disentanglement in the background will benefit from further work, since it is comprised of both technical artifacts and biological variations. This can be addressed in future work by extending our model to include three encoders corresponding to three latent spaces that separately captures technical noise, natural (biological) phenotypic variations, and novel perturbation-induced phenotypes. We can incorporate kernel-based independence measures [19] to facilitate the enforcement of independence statements between the technical noise latent variables and the perturbation label.

Exploring deeper neural network architectures is another important extension of this work. In this current work, we used a relatively simple encoder/decoder architecture with only five convolutional/deconvolutional layers. A deeper architecture might foster a finer granularity in the detection of subtle phenotypic patterns that are otherwise overshadowed in shallow architectures. Furthermore, we used a small number of dimensions for the salient and background space (32 dimensions each). Increasing the number of latent dimensions in our mcVAE model can potentially enhance the representation of complex, high-dimensional data, allowing for a more nuanced understanding of genetic perturbations.

## A. Supplementary Material

### A.1 Data acquisition and preprocessing

CellProfiler features with corresponding metadata, including cell cycle stage and perturbation label (gene targeted by CRISPR), for each cell were obtained directly from the online repository Harvard Dataverse [20]. These features were already batch-corrected by standardizing against the NTCs in the corresponding batch.

Raw microscopy images each covering a large field of view with many cells were downloaded from BioImage Archive [21], followed by imaging channel alignment using phase cross-correlation. For each raw image, pixels with intensity values in the top and bottom 0.1% were clipped. Finally, individual cell patches were obtained by using the cell positional values from CellProfiler data to obtain a 100 pixel by 100 pixel bounding box around each cell, which was used to represent individual cells for model training. We removed cells within 50 pixel of the tile edge.

## Acknowledgments and Disclosure of Funding

We thank Avtar Singh, David Richmond, Mahtab Bigverdi, Anqi Zhu, Rebecca Boiarsky, Xinming Tu, and Kexin Huang for insightful feedback throughout the duration of this project which greatly improved this work. We also thank members of the Regev Lab and the Artificial Intelligence/Machine Learning department at Genentech for providing constructive feedback on the results presented in this work.

## Code Availability Statement

Multi-Contrastive VAE is implemented in PyTorch, with all implementation code for model training, analysis, and figure generation available at https://github.com/Genentech/contrastive-ops.

